# Molecular Drivers of Tumor Progression in Microsatellite Stable *APC* Mutation-Negative Colorectal Cancers

**DOI:** 10.1101/2021.09.14.460319

**Authors:** Adam Grant, Rosa M. Xicola, Vivian Nguyen, James Lim, Curtis Thorne, Bodour Salhia, Xavier Llor, Nathan Ellis, Megha Padi

## Abstract

**Background:** The tumor suppressor gene adenomatous polyposis coli (*APC*) is the initiating mutation in approximately 80% of all colorectal cancers (CRC), underscoring the importance of aberrant regulation of intracellular WNT signaling in CRC development. Recent studies have found that early-onset CRC exhibits an increased proportion of tumors lacking an *APC* mutation. We set out to identify mechanisms underlying *APC* mutation-negative (*APC*^*mut–*^) CRCs.

**Methods:** We analyzed data from The Cancer Genome Atlas to compare clinical phenotypes, somatic mutations, copy number variations, gene fusions, RNA expression, and DNA methylation profiles between *APC*^*mut–*^ and *APC* mutation-positive (*APC*^*mut+*^) microsatellite stable CRCs.

**Results:** Transcriptionally, *APC*^*mut–*^ CRCs clustered into two approximately equal groups. Cluster One was associated with enhanced mitochondrial activation. Cluster Two was strikingly associated with genetic inactivation or decreased RNA expression of the WNT antagonist *RNF43*, increased expression of the WNT agonist *RSPO3*, activating mutation of *BRAF*, or increased methylation and decreased expression of *AXIN2. APC*^*mut–*^ CRCs exhibited evidence of increased immune cell infiltration, with significant correlation between M2 macrophages and *RSPO3*.

**Conclusions:** *APC*^*mut–*^ CRCs comprise two groups of tumors characterized by enhanced mitochondrial activation or increased sensitivity to extracellular WNT, suggesting that they could be respectively susceptible to inhibition of these pathways.

## BACKGROUND

Colorectal cancer (CRC) is the second deadliest cancer in the United States, with an estimated 147,950 individuals diagnosed and 53,200 deaths in 2020^1^. Although there have been great reductions in CRC incidence and mortality widely attributed to increased screening^2^, the incidence of CRC has been increasing in individuals less than 50 years of age at a rate of 2% per year since 1994^3^. Molecular analysis has shown that <20% of early-onset CRC cases are explained by genetically determined hereditary syndromes^4^ and a variety of environmental factors have been postulated to underlie its increase^5^, suggesting that a unitary cause of early-onset CRC will be elusive. With early-onset CRC manifesting as a heterogenous disease caused by a multitude of factors, there is a pressing need to identify the distinct molecular subtypes of CRC that are overrepresented in early-onset cases.

Somatic mutation of the adenomatous polyposis coli (*APC*) gene is the initiating event in approximately 80% of all CRCs, but *APC* mutations are significantly less frequent in early-onset CRCs^6-8^. *APC* is a structural and regulatory component of a destruction complex, which responds to WNT stimulation by inhibition of degradation of the stem cell-promoting transcription factor β-catenin, encoded by the *CTNNB1* gene^9^. Failure to regulate β-catenin by degradation due to mutational inactivation of *APC* throws colorectal epithelial cells into a continuous “WNT-activated” state; they no longer require activation by WNTs to maintain the stem cell compartment^10^. The fact that early-onset CRCs more frequently lack an *APC* mutation suggests that many of these tumors depend on alternative molecular mechanisms. In mismatch repair-deficient CRCs, which exhibit microsatellite instability (MSI) and constitute 12-15% of all CRCs, *APC* mutations are also significantly less frequent and *BRAF* mutations constitute a dominant driver mechanism^11^. What initiates and drive the carcinogenetic process in microsatellite stable (MSS) CRCs that lack *APC* mutations? Here, we comprehensively compare molecular profiles of MSS *APC* mutation-positive CRCs (*APC*^*mut+*^) and MSS *APC* mutation-negative (*APC*^*mut–*^) CRCs to identify novel *APC*-independent mechanisms driving CRC subtypes.

## METHODS

### Analyses of genomic alterations in case series

To formulate a discovery series, we obtained colon adenocarcinoma (COAD) and rectal adenocarcinoma (READ) Mutect2 MAF files from The Cancer Genome Atlas (TCGA) data in the Genomic Data Commons portal (Supplementary File 1). Curated somatic nucleotide variant and copy number data were extracted using TCGAbiolinks and FireBrowse, respectively^12^. Deep deletions, amplifications, and gene fusions^13^ were identified. We excluded hypermutable cases by removing MSI-high cases based on clinical data and by removing cases with >700 mutations. CRC samples were classified as *APC*^*mut–*^ if they lacked a non-silent or deep mutation in *APC* or lacked a mutation in *CTNNB1*^14^. With these filtration steps, we had 63 *APC*^*mut–*^ samples and 362 *APC*^*mut+*^ samples. We compared genomic alterations between *APC*^*mut–*^ and *APC*^*mut+*^ CRCs by Fisher’s exact test and tested mutual exclusivity by CoMEt^15^. For more details on the bioinformatics analysis, see Supplementary Methods.

For validation series, we used the CPTAC-2^16^ and GSE35896^17^ datasets, because they were the only CRC datasets with *APC* mutation status and gene expression data available from the cBioPortal, the International Cancer Genome Consortium or studies utilized by Guinney *et al*. to determine consensus molecular subtypes of CRC^18^. CPTAC-2 was downloaded from cBioPortal and GSE3896 from synapse.org^18^. In the CPTAC-2 dataset, we identified 11 *APC*^*mut–*^ CRCs and 70 *APC*^*mut+*^ CRCs. Because GSE35896 did not have whole exome sequencing or copy number data, we could not filter for hypermutation, *APC* deep deletion, or *CTNNB1* mutations. Based on the data available, we classified 16 out of 56 MSS CRC samples as *APC*^*mut–*^.

### Transcriptomic analyses

For TCGA and CPTAC-2 series, we obtained HTSeq count files and used the edgeR and limma pipeline to normalize the counts matrix. For GSE35896, we used the RMA normalized data. Genes that had low CPMs were discarded. MBatch analysis showed no evidence of batch effects.

We used limma to identify differentially expressed genes between *APC*^*mut–*^ and *APC*^*mut+*^ CRCs (P_adj_ < 0.05). PathView was used to map differentially expressed genes onto the KEGG WNT canonical signaling pathway, with the node sum parameter set to “max.abs”^19^. Gene set enrichment analysis (GSEA) was performed using fgsea^20^. For GSEA input, we used the c5.go.bp.v7.4.symbols.gmt.txt file from msigdb.org and ranked the genes by the t-statistic from our differential expression analysis. The Cytoscape application EnrichmentMap was used to visualize all statistically significant GSEA GO terms (P_adj_ < 0.05)^21^. CIBERSORTx was used to impute the fraction of immune cells based on gene expression data from the TCGA, GSE35896, and CPTAC-2 datasets^22^.

To quantify the WNT signal-transduction competence, we defined the WNT ligand sensitivity (WNT_LS_) score as the normalized expression of *RSPO3* minus the sum of the normalized expression levels of *RNF43* and *ZNRF3*,

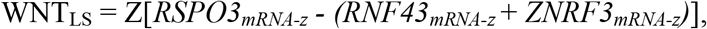

where the subscript mRNA-z denotes the Z-score of the expression value relative to all samples including tumors and normals. The WNT_LS_ score for each tumor was then compared to the maximum WNT ligand expression over all 12 WNTs in the same tumor.

### DNA methylation analyses

IDAT files for the TCGA Illumina Human Methylation 450 arrays were retrieved using GDCquery from TCGAbiolinks. Preprocessing and normalization were carried out with the preprocessFunnorm method in minfi^23^. MBatch analysis showed no evidence of batch effects. Differentially methylated regions (DMRs) were identified using DMRcate and annotated with annotatr^24,25^. For DMRs that spanned multiple gene regions, we selected the gene with the most significant beta-values. To quantify methylation of a DMR, we took the average of all the statistically significant beta-values associated with the DMR.

### DepMap data analyses

DepMap data was obtained from https://depmap.org/portal/download/^26^. CRC cell lines were selected excluding those with MSI and with >800 mutations. To distinguish *APC*^*mut–*^ from *APC*^*mut+*^ CRC cell lines, we used the protocol for the TCGA dataset. To rank CRISPR knockouts, we used the Welch’s two-sample t-test statistic applied to the dependency scores of *APC*^*mut–*^ and *APC*^*mut+*^ cell lines. Dependency scores were extracted from the file “Achilles_gene_dependency.csv” on the DepMap portal.

#### Ethics statement

Ethics approval is not required for this study because it does not involve human participants or animal subjects.

## RESULTS

### Age effect in *APC*^*mut–*^ CRCs

To identify characteristics that distinguished *APC*^*mut–*^ from *APC*^*mut+*^ CRCs, we compared molecular profiles between the two groups in a discovery series from the TCGA, then validated the results in two additional publicly available series. CRC samples that exhibited MSI or were hypermutated were excluded from our study, because tumors with these characteristics constitute a well-defined subtype^11^. In addition to separating MSS and non-hypermutated CRCs by *APC* mutation status, samples that contained a *CTNNB1* mutation^14^ or deep deletion of *APC* were also classified as *APC*^*mut+*^. After applying these filtration steps, we classified 63 of 425 (15%) of the MSS CRCs in the TCGA dataset as *APC*^*mut–*^. In the GSE35896 validation dataset, 16 out of 56 (29%) CRCs were classified as *APC*^*mut–*^ and in the CPTAC-2 dataset, 11 out of 81 (14%) CRCs were classified as *APC*^*mut–*^.

We tested clinical features that might be statistically associated with TCGA *APC*^*mut–*^ CRCs (Table 1). In agreement with previous studies^6-8^, *APC*^*mut–*^ CRCs were diagnosed at a younger age (61.4 in *APC*^*mut–*^ vs. 66.4 in *APC*^*mut+*^), and 63% of tumors diagnosed <50 were *APC*^*mut–*^. *APC*^*mut–*^ CRCs were also younger in the CPTAC-2 dataset (61.5 in *APC*^*mut–*^ vs. 65.5 in *APC*^*mut+*^), but this observation did not reach statistical significance (p = 0.24). (Age of diagnosis was not available for the GSE35896 dataset.) In addition to age, TCGA *APC*^*mut–*^ CRCs were more prevalent in Asians (p = 0.005), were more likely to be classified as CpG island methylator phenotype (CIMP) high (p = 0.02), and were more likely to be diagnosed later than stage one (p = 0.035).

**Table 1.** Comparison of clinical in *APC* mutation-positive (*APC*^*mut+*^) and *APC* mutation-negative (*APC*^*mut–*^) colorectal cancers. P-values were calculated for comparisons between *APC*^*mut+*^ CRCs and *APC*^*mut–*^ CRCs from the TCGA dataset. A two-sample t-test with a two-tailed p-value was performed for continuous features and a Fisher’s exact test with a two-tailed p-value was performed for categorical data. A p-value threshold of 0.05 was considered significant. CIMP, CpG island methylator phenotype was defined by unsupervised clustering as reported by Guinney *et al*. CMS, consensus molecular subtypes of colorectal cancer determined by Guinney *et al*.

| Feature | *APC*<sup>mut+</sup><br>(N = 362, 85%) | *APC*<sup>mut-</sup><br>(N = 63, 15%) | P-Value |
| --- | --- | --- | --- |
| Age | 66.4 | 61.4 | **.004** |
| Non-silent Mutations | 121.4 | 112.4 | .21 |
| Male/Female | 194/165 (54%) | 30/33 (48%) | .41 |
| COAD/READ | 250/109 (70%) | 51/12 (81%) | .07 |
| Proximal/Distal | 104/195 (35%) | 24/25 (49%) | .08 |
| White | 305/359 (85%) | 53/63 (84%) | .85 |
| African American | 46/359 (13%) | 5/63 (8%) | .40 |
| Asian | 4/359 (1%) | 5/63 (8%) | **.005** |
| American Indian | 4/359 (1%) | 0/63 (0%) | 1.0 |
| Stage I | 62/346 (18%) | 4/59 (7%) | **.035** |
| Stage II | 106/346 (31%) | 23/59 (39%) | .23 |
| Stage III | 119/346 (34%) | 18/59 (31%) | .66 |
| Stage IV | 59/346 (17%) | 14/59 (23%) | .27 |
| CIMP-0 | 177/251 (70%) | 30/50 (60%) | .18 |
| CIMP-Low | 57/251 (23%) | 11/50 (22%) | 1.0 |
| CIMP-High | 17/234 (7%) | 9/50 (18%) | **.02** |
| CMS1 | 5/291 (2%) | 2/49 (4%) | .27 |
| CMS2 | 149/291 (51%) | 20/49 (41%) | .22 |
| CMS3 | 41/290 (14%) | 7/49 (14%) | 1.0 |
| CMS4 | 96/290 (33%) | 20/49 (41%) | .33 |

### WNT signaling mutations in *APC*^*mut–*^ CRCs

To identify distinctive somatic mutations, we compared non-silent nucleotide variants, gene amplifications, deep gene deletions, and gene fusions in *APC*^*mut–*^ and *APC*^*mut+*^ CRCs (Figure 1A). The top three most statistically different genomic alterations specific to *APC*^*mut–*^ CRCs were *PTPRK-RSPO3* gene fusions (p = 1.3 × 10^−5^), *RNF43* mutations (p = 4.7 × 10^−5^) and *BRAF* mutations (p = 1.9 × 10^−4^). These genetic alterations have been identified in CRC previously with evidence for mutual exclusivity with *APC* mutations^27-30^. (The *RNF43* mutation G659Vfs*41, which is associated with MSI CRCs, was not present in the tumors analyzed here^31^.) Eight of nine *BRAF* mutated *APC*^*mut–*^ CRCs had the oncogenic V600E *BRAF* mutation. Six of eight *RNF43* mutated *APC*^*mut–*^ CRCs had mutations that caused premature protein truncations, whilst one sample had a previously identified missense mutation, R554G. These findings suggested that *BRAF* and *RNF43* mutations are associated with tumor progression in MSS *APC*^*mut–*^ CRCs.

**Figure 1.**
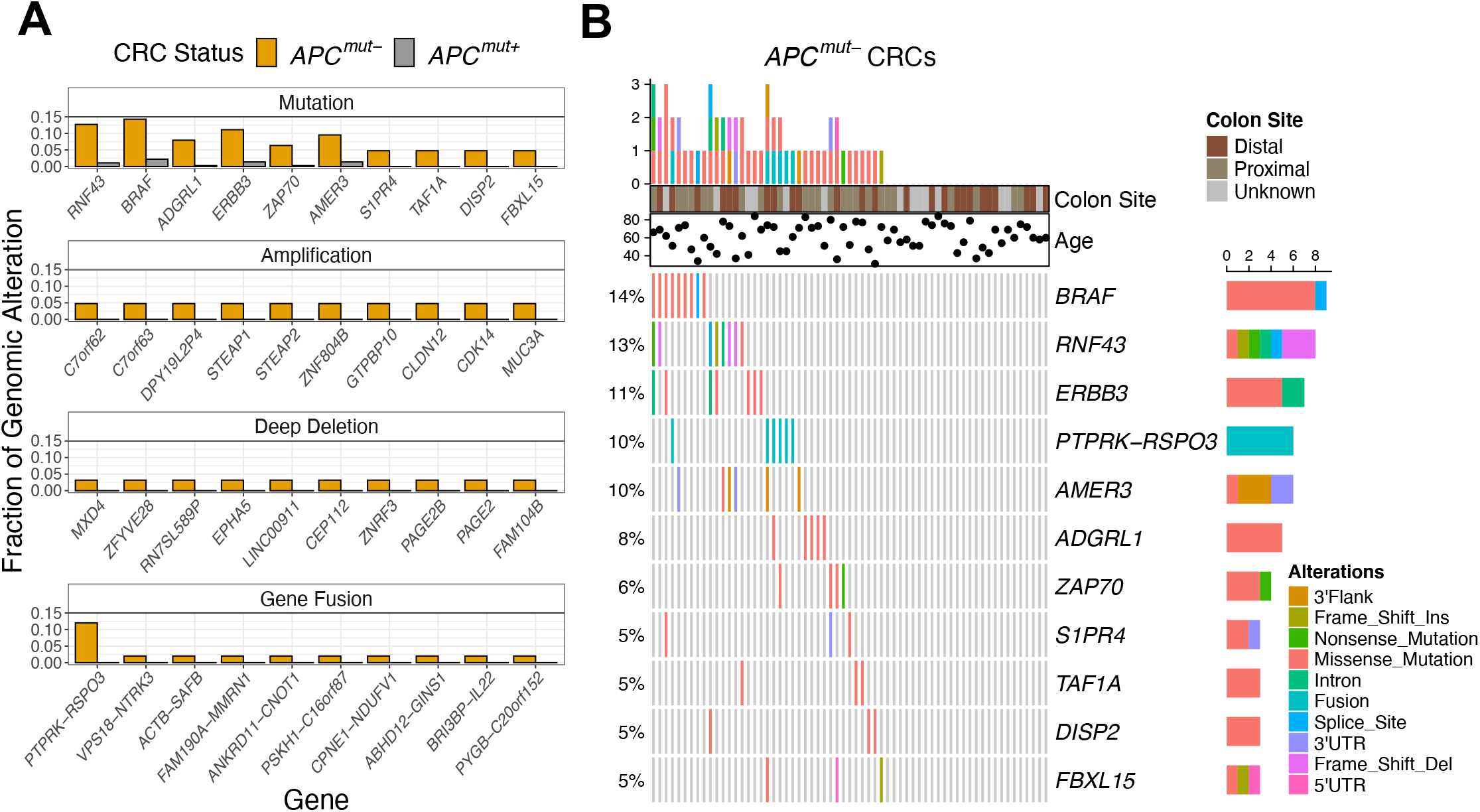
WNT signaling mutations in *APC*^*mut–*^ CRCs. (*A*) Fraction of *APC*^*mut–*^ CRCs from the TCGA dataset with gene mutations, amplifications, deep deletions, and fusions that were significantly more common in *APC*^*mut–*^ in comparison to *APC*^*mut+*^ CRCs. The top 10 are shown by p-value ranking, most significant (left) to least (right). (*B*) OncoPrint diagram showing the top 10 statistically significant mutations associated with *APC*^*mut–*^ CRCs and the gene fusion *PTPRK-RSPO3* for the 63 *APC*^*mut–*^ CRCs.

Based on the mutated genes in Figure 1A and the *PTPRK-RSPO3* gene fusion, we found that only 37 out of 63 samples (59%) contained a genomic alteration that distinguished *APC*^*mut–*^ from *APC*^*mut+*^ CRCs (Figure 1B). No pairwise combination of genes were statistically mutually exclusive. However, *PTPRK-RSPO3* gene fusions and *RNF43* mutations never co-occurred and were found in 23% of the *APC*^*mut–*^ CRCs. After disregarding overlapping genomic alterations, *BRAF* mutations were the next most abundant (10%), followed by mutations in *ADGRL1* (6%), *ERBB3* (5%), and *ZAP70* (5%). Supporting these findings, we found that *BRAF* mutations in the GSE35896 dataset and mutations in *RNF43, ERBB3*, and *ZAP70* in the CPTAC-2 dataset were more frequent in *APC*^*mut–*^ CRCs than in *APC*^*mut+*^ CRCs (Supplementary Figure 1). (No additional mutation information was provided with the GSE35896 dataset.)

### Enhanced sensitivity to extracellular WNT in *APC*^*mut–*^ CRCs

Because a distinctive somatic mutational mechanism was not evident in over 40% of *APC*^*mut–*^ CRCs, we examined transcriptomics data for further distinguishing molecular characteristics. Strikingly, in differential gene expression analysis of the TCGA dataset, *RNF43* was the most differentially expressed gene between the two tumor groups (P_adj_ = 4.6 ⨯ 10^−15^; Figure 2A), with a -0.98 log_2_ fold decrease in mean expression level in *APC*^*mut–*^ CRCs. Consistent with these results, *RNF43* was also down-regulated in *APC*^*mut–*^ CRCs in the GSE35896 and CPTAC-2 validation datasets (Figure 2B). *RNF43* and its family member *ZNRF3* are membrane-bound E3 ubiquitin ligases that actuate the degradation of low-density-lipoprotein-related protein (LRP)-FZD WNT receptors. They associate with leucine-rich repeat-containing G-protein coupled receptors (LGR) that down-regulate the levels of *RNF43* and *ZNRF3* at the surface upon binding of R-SPONDINs^32^. The transcriptional down-regulation of *RNF43* suggested that *APC*^*mut–*^ CRCs may express higher levels of LRP-FZD receptors at the cell surface, and consequently be more responsive to extracellular WNTs.

**Figure 2.**
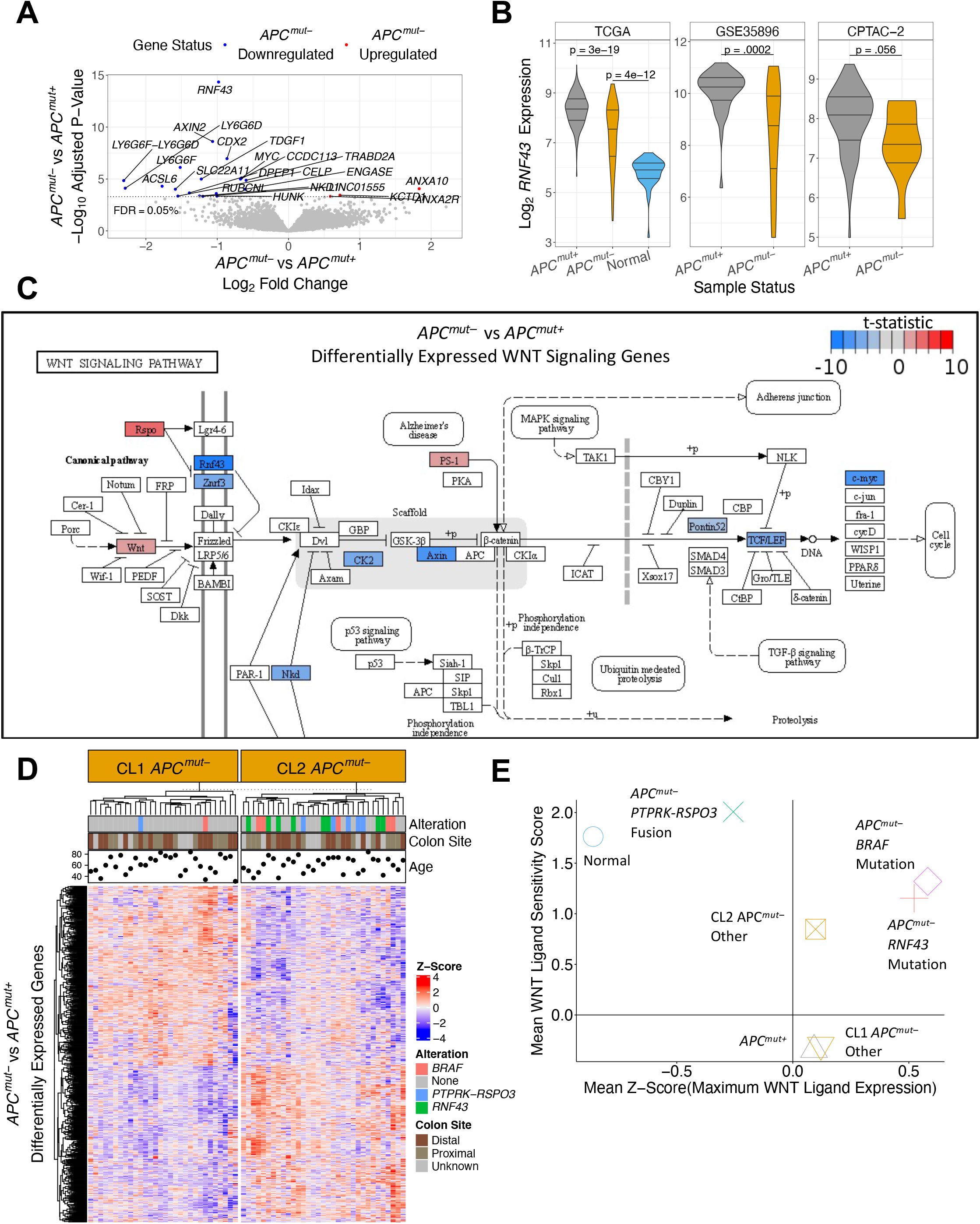
Enhanced sensitivity to extracellular WNT in *APC*^*mut–*^ CRCs. (*A*) Volcano plot representing the results from differential expression analysis between *APC*^*mut–*^ and *APC*^*mut+*^ CRCs. Labeled points are the genes with an P_adj_ < 0.0005. Blue points were downregulated in *APC*^*mut–*^ CRCs and red points upregulated. (*B*) Comparison of *RNF43* gene expression in *APC*^*mut–*^ CRCs, *APC*^*mut+*^ CRCs, and normal colon samples in the TCGA, GSE35896 and CPTAC-2 datasets. Two-sample t-tests with a two tailed p-value were used to test statistical significance. (*C*) Differentially expressed genes (P_adj_ < 0.05) between *APC*^*mut–*^ and *APC*^*mut+*^ CRCs from TCGA were mapped onto the KEGG canonical WNT signaling pathway. Blue labeling represents genes downregulated in *APC*^*mut–*^; red labeling represents upregulated genes. (*D*) Unsupervised clustering analysis of *APC*^*mut–*^ CRCs from the TCGA dataset using differentially expressed genes (P_adj_ < 0.05). (*E*) Scatter plot showing estimation of activation potential of extracellular WNT signaling. Each point is the mean for individual groups. The y-axis represents a group’s apparent sensitivity to extracellular WNT signaling using the WNT ligand sensitivity score. The x-axis represents a group’s WNT stimulation potential by quantifying each sample’s maximum WNT ligand expression.

Because WNT signaling was implicated by these results, we sought to determine the extent to which other factors in the canonical WNT signaling pathway were differentially expressed between *APC*^*mut–*^ and *APC*^*mut+*^ CRCs (Figure 2C). Consistent with the results above, we observed that other genes involved in extracellular WNT signaling were dysregulated, namely *RSPO3* and *ZNRF3*. Differences in *RSPO3* and *ZNRF3* mRNA expression showed a similar trend in the validation datasets and were statistically significant in select cases (Supplementary Figure 2). We did not observe any differential expression of the extracellular WNT regulator genes *LGR4, LGR5, LGR6*, or LRP-FZD receptors. The fact that *LGR4-6* were not differentially expressed between *APC*^*mut–*^ and *APC*^*mut+*^ CRCs was consistent with the finding that *RSPO3* does not require interaction with LGRs to potentiate WNT signaling^33^ and LRP-FZD receptor levels are regulated post-transcriptionally^34^. When we compared gene expression of *APC*^*mut–*^ and *APC*^*mut+*^ CRCs to normal samples and mapped genes onto the canonical WNT signaling pathway, changes in gene expression in WNT signaling were similar between these two tumor types (Supplementary Figure 3). These results suggested that both types of CRCs exploit changes in WNT signaling. However, based on the mutation and expression data, *APC*^*mut–*^ CRCs appear to favor dysregulation of genes involved in response to extracellular WNT signaling, whereas *APC*^*mut+*^ CRCs are stuck in the “on” state and are WNT signal-transduction incompetent.

To determine the fraction of *APC*^*mut–*^ CRCs that operate via enhanced sensitivity of extra-cellular WNT, we performed unsupervised hierarchical clustering using all differentially expressed genes (P_adj_ < 0.05) between *APC*^*mut–*^ and *APC*^*mut+*^ CRCs in the TCGA dataset (Figure 2D). *APC*^*mut–*^ CRCs clustered into two prominent groups, referred to here as Cluster 1 (CL1) and Cluster 2 (CL2). Most *APC*^*mut–*^ CRCs with a *PTPRK-RSPO3* fusion, *BRAF* mutation, or *RNF43* mutation were in CL2. To estimate response to extracellular WNT signaling, we developed a WNT ligand sensitivity (WNT_LS_) score and compared it to maximum WNT ligand expression (Figure 2E; see Methods for more details). We postulated that CRCs that have a high WNT_LS_ score and high WNT ligand expression are driven by extracellular WNT signaling. Consistent with this logic, *APC*^*mut–*^ CRCs with *RNF43* mutations had higher WNT_LS_ scores than *APC*^*mut+*^ CRCs and higher maximum WNT expression than normals. Interestingly, *APC*^*mut–*^ CRCs with *PTPRK-RSPO3* fusions were the most sensitive to WNT ligands, but had the lowest expression of WNT ligands compared to other CRCs. Inconsistencies in how CRCs with *PTPRK-RSPO3* fusions and CRCs with *RNF43* mutations enhance their sensitivity to extracellular WNT signaling may be due to different evolutionary pressures.

*APC*^*mut–*^ CRCs with *BRAF* mutations also exhibited higher WNT_LS_ and higher WNT ligand expression, similar to *APC*^*mut–*^ CRCs with *RNF43* mutations. Importantly, *APC*^*mut–*^ CRCs from CL2 that did not have *BRAF* mutations, *RNF43* mutations, or *PTPRK-RSPO3* fusions exhibited higher WNT_LS_ scores compared to *APC*^*mut+*^ CRCs. In contrast, *APC*^*mut–*^ CRCs from CL1 exhibited WNT_LS_ scores similar to *APC*^*mut+*^ CRCs. *APC*^*mut–*^ CRCs from the GSE35896 and CPTAC-2 datasets also clustered into two groups with high and low WNT_LS_ scores (Supplementary Figure 4). Given the importance of WNT signaling in CRC, these results suggest that other WNT-related mechanisms drive CL1 *APC*^*mut–*^ CRCs. By transcriptomic analysis, CL1 *APC*^*mut–*^ CRCs were practically indistinguishable from *APC*^*mut+*^ CRCs; however, GSEA showed enrichment of oxidative phosphorylation genes (Supplementary Figure 5 and 6), implicating mitochondrial activation in CL1 *APC*^*mut–*^ tumorigenesis. These data were supported by data from the DepMap CRISPR screen that demonstrated dependence of *APC*^*mut–*^ CRC cell lines on oxidative phosphorylation complexes in the mitochondria (Supplementary Figure 5E).

### *APC*^*mut–*^ CRCs associated with immune infiltration

GSEA analysis showed that GO terms related to the adaptive immune response were upregulated in *APC*^*mut–*^ compared to *APC*^*mut+*^ CRCs (Figure 3A). To further investigate immune system involvement in *APC*^*mut–*^ CRCs, we employed the bulk tissue deconvolution method CIBERSORTx^22^. In agreement with the GSEA results, the CIBERSORTx absolute score was increased in *APC*^*mut–*^ compared to *APC*^*mut+*^ CRCs in all three CRC datasets (Figure 3B). The CIBERSORTx absolute score was highest in *APC*^*mut–*^ CRCs with *BRAF* or *RNF43* mutations and CL2 *APC*^*mut–*^ CRCs without mutations (Figure 3C). Because these *APC*^*mut–*^ CRCs had more infiltrating immune cells than those with *PTPRK-RSPO3* fusions, we tested whether any of the 22 immune cell types were associated with expression of WNT agonist ligand *RSPO3* (Figure 3D). We found that M2 macrophages had the strongest positive Pearson correlation with *RSPO3* expression. M2 macrophages and *RSPO3* expression were also significantly correlated in the GSE35896 and the CPTAC-2 datasets. Macrophage expression of *RSPO3* was shown in a study of patients with pulmonary fibrosis^35^.

**Figure 3.**
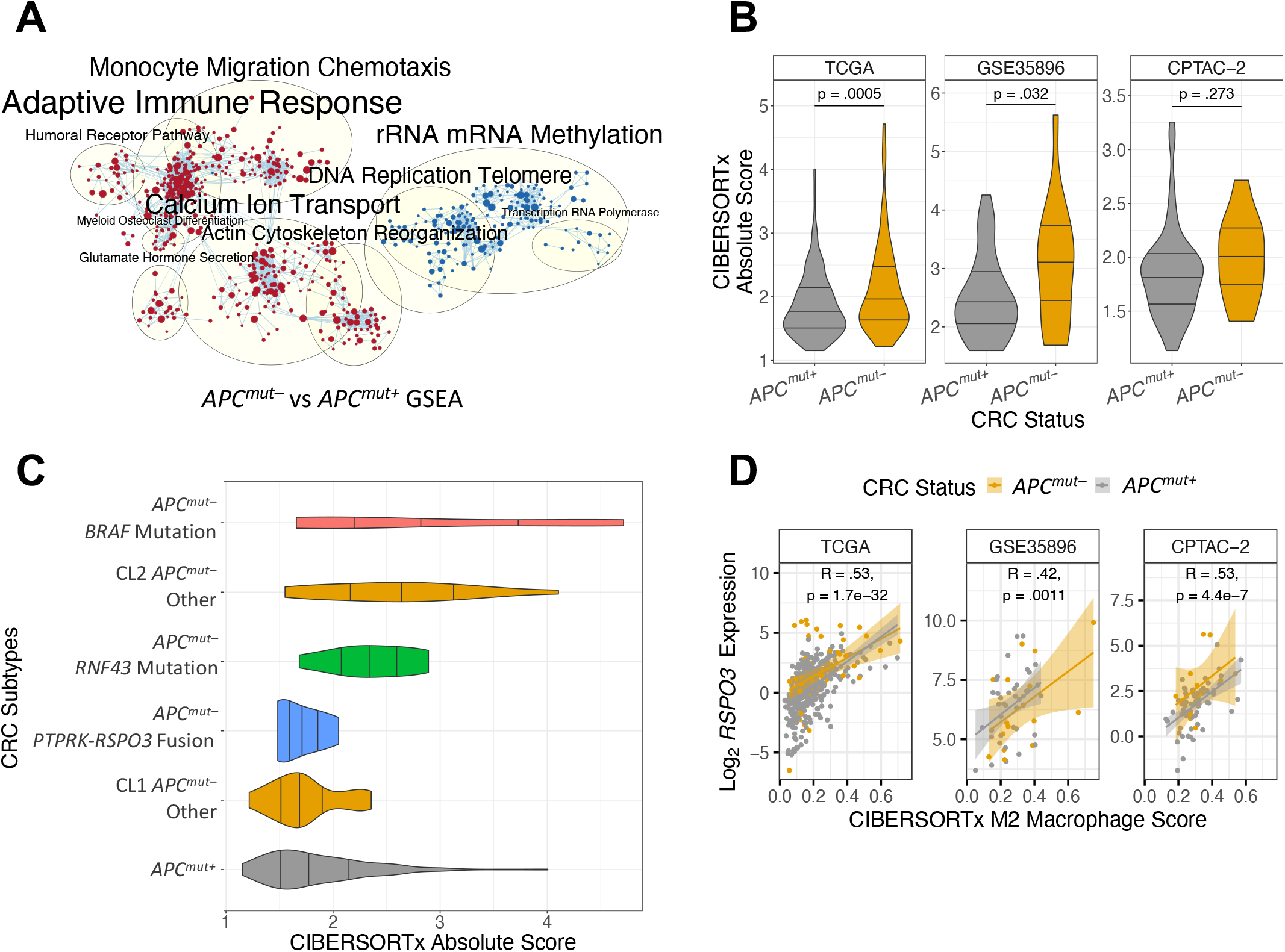
*APC*^*mut–*^ CRCs associated with immune infiltration. (*A*) GSEA results of differential gene expression analysis of *APC*^*mut–*^ vs *APC*^*mut+*^ CRCs from the TCGA dataset. Red clusters represent GO terms enriched among upregulated genes in *APC*^*mut–*^ CRCs and blue clusters correspond to down-regulated processes. (*B*) CIBERSORTx absolute score in CRCs from the TCGA, GSE35896 and CPTAC-2 datasets. Two-sample t-tests with a two-tailed p-value were used to test statistical significance. (*C*) Violin plot of CIBERSORTx absolute score across subtypes of *APC*^*mut–*^ CRCs. (*D*) Expression of *RSPO3* in *APC*^*mut–*^ and *APC*^*mut+*^ CRCs plotted against their individual M2 macrophage scores identified from the CIBERSORTx algorithm. Pearson correlation was performed to determine statistical significance.

### *APC*^*mut–*^ CRCs have higher *AXIN2* methylation

Because we found an association between *APC*^*mut–*^ CRCs and CIMP-high previously^7^, we identified differentially methylated regions (DMRs) between *APC*^*mut–*^ and *APC*^*mut+*^ CRCs. *APC*^*mut–*^ CRCs were globally more hypermethylated than *APC*^*mut+*^ CRCs, with a particular excess in promoter regions (Figure 4A). Comparing the top ten hypermethylated and hypomethylated DMRs, we did not observe the same statistically significant genes as we did in the mutation and expression analyses (Figure 4B). However, when we tested correlation of *RNF43* expression with DNA methylation levels of DMRs and with RNA expression, we found that methylation and gene expression of *AXIN2* had the highest correlations (Figure 4C). *RNF43* gene expression was also significantly correlated with *AXIN2* expression in the GSE35896 and CPTAC-2 datasets (Figure 4D; methylation data was not available in these datasets). Increased *AXIN2* DNA methylation was associated with decreased *RNF43* expression in a subset of *APC*^*mut–*^ CRCs that did not have one of the common somatic mutations (Figure 4E). Similar to our findings with *RSPO3* expression, we found that M2 macrophages correlated most with *AXIN2* DNA methylation (Figure 4F).

**Figure 4.**
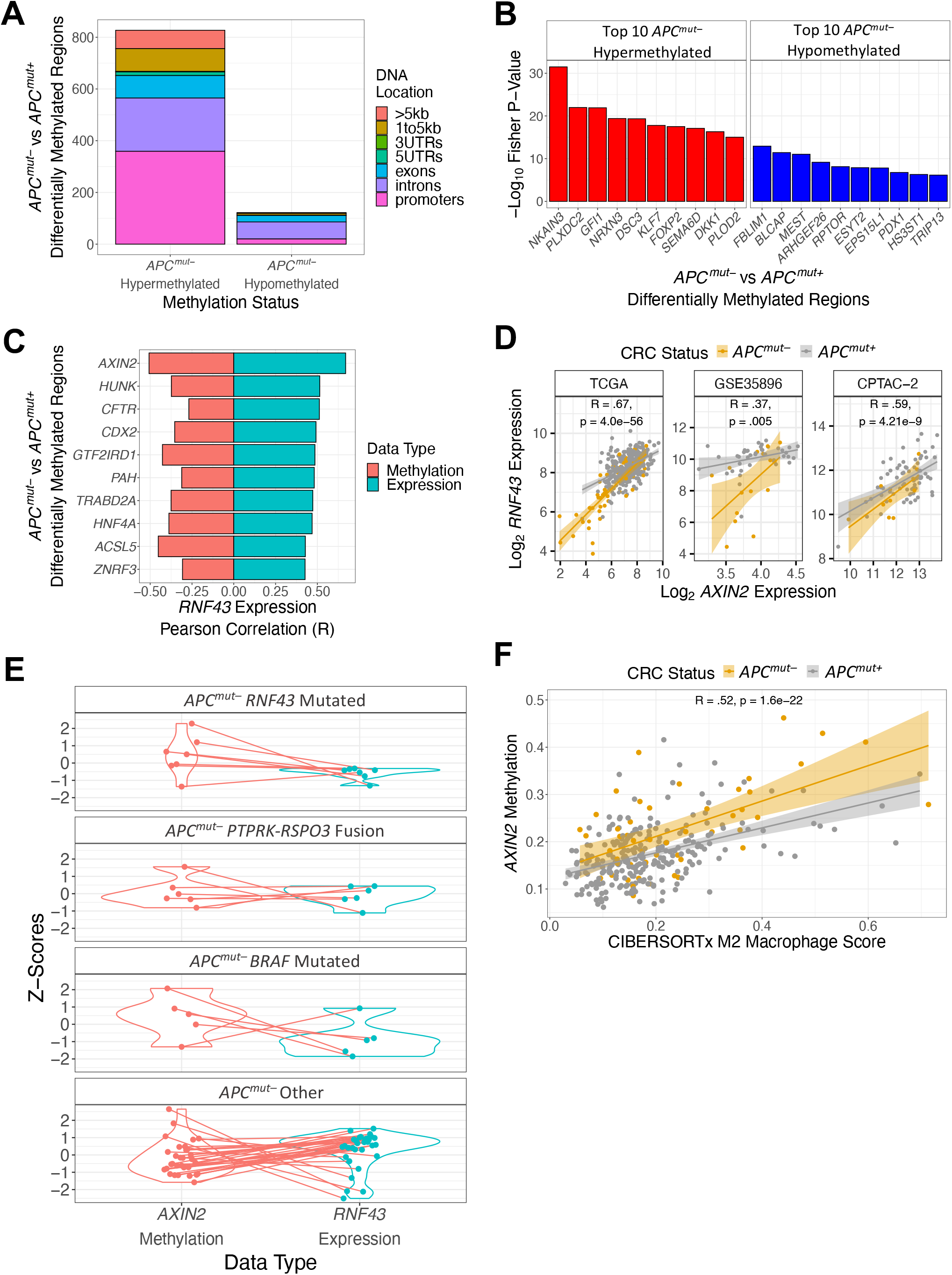
*APC*^*mut–*^ CRCs have higher *AXIN2* methylation. (*A*) Bar plot comparing total number of hyper-methylated and hypo-methylated differentially methylated regions (DMRs) between *APC*^*mut–*^ and *APC*^*mut+*^ CRCs from the TCGA dataset. (*B*) Top 10 *APC*^*mut–*^ hypermethylated and hypomethylated DMRs between *APC*^*mut–*^ and *APC*^*mut+*^ CRCs from TCGA. Red bars represent *APC*^*mut–*^ CRC differentially hypermethylated genes and blue bars represent *APC*^*mut–*^ CRC differentially hypomethylated genes. (*C*) Bar plot representing DMRs with strongest correlations with *RNF43*. Blue bars represent the top 10 DMRs with the highest Pearson gene expression correlation with *RNF43* gene expression. Red bars represent the Pearson correlation between the average differentially methylated beta values and *RNF43* expression for these differentially methylated regions. (*D*) Scatter plots of *RNF43* expression and *AXIN2* expression of both *APC*^*mut–*^ and *APC*^*mut+*^ CRCs in the TCGA, GSE35896, and CPTAC-2 datasets. Pearson correlation was performed to determine statistical significance. (*E*) Matched comparison between Z-normalized *AXIN2* average beta values and Z-normalized *RNF43* expression of *APC*^*mut–*^ CRCs. (*F*) Scatter plot of *AXIN2* average beta values and the CIBERSORTx M2 macrophage score of *APC*^*mut–*^ and *APC*^*mut+*^ CRCs from the TCGA dataset. Pearson correlation was performed to measure statistical significance.

### *AP2M1* gene expression associated with earlier onset in *APC*^*mut–*^ CRCs

Age of onset was not different in CL1 and CL2 *APC*^*mut–*^ CRCs (Figure 5A). To identify gene expression changes linked to earlier age of onset in *APC*^*mut–*^ CRCs, we separated *APC*^*mut–*^ CRCs into two groups based on the median expression of each gene and performed a logrank test between these two groups, using the age at diagnosis as the event variable. Expression of *AP2M1* best distinguished earlier onset *APC*^*mut–*^ CRCs from later onset *APC*^*mut–*^ CRCs (Figure 5B), and higher *AP2M1* expression was associated with earlier onset in *APC*^*mut–*^ relative to *APC*^*mut+*^ CRCs (Figure 5C).

**Figure 5.**
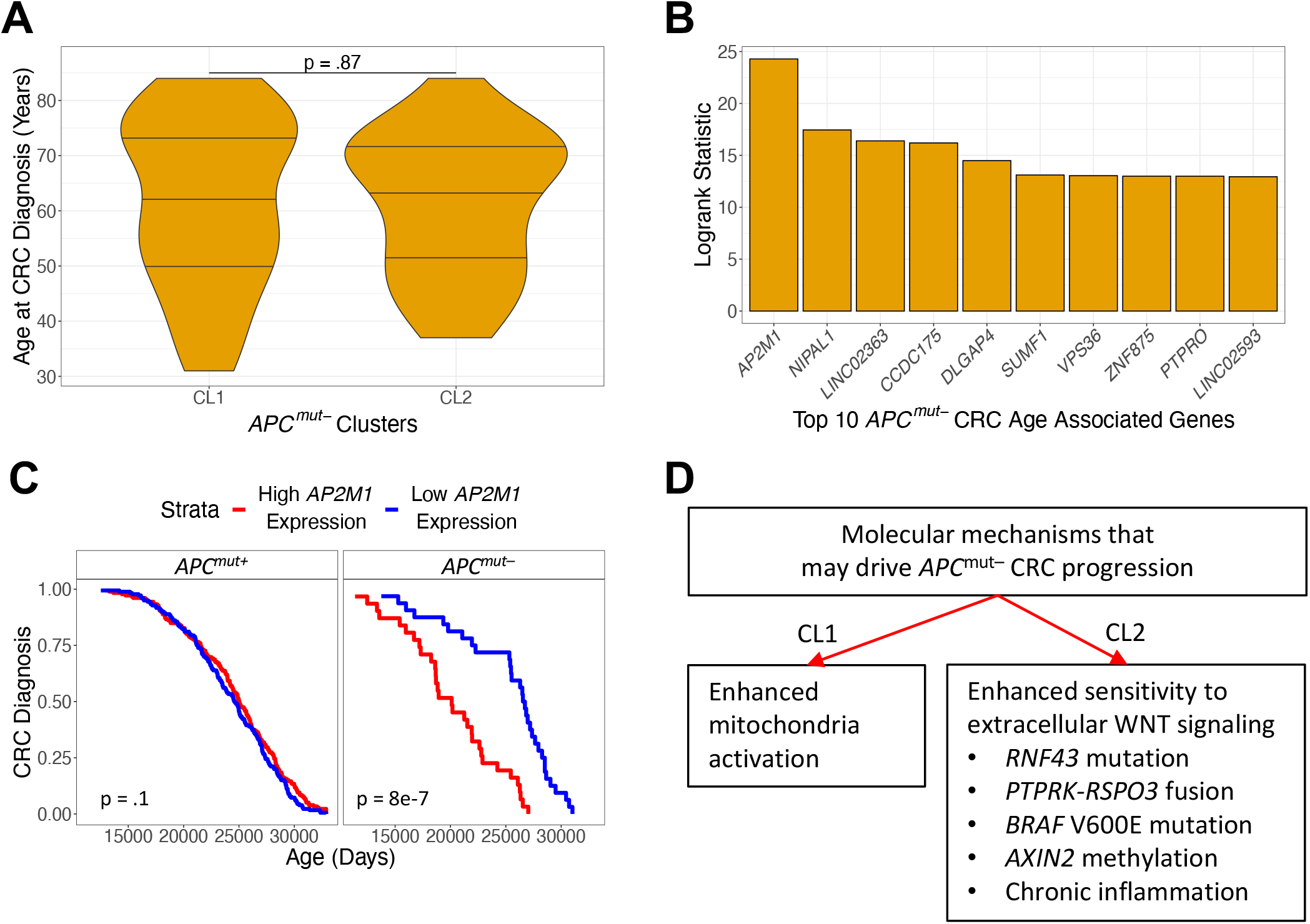
*AP2M1* gene expression associated with earlier-onset in *APC*^*mut–*^ CRCs. *(A)* A comparison of age between *APC*^*mut–*^ clusters identified from Figure 2D. A two-sample t-test with a two-tailed p-value was used to determine statistical significance. (*B*) Top 10 statistically significant genes based on a logrank test whose median gene expression best separates age of CRC diagnosis of *APC*^*mut–*^ CRCs from the TCGA dataset. (*C*) Kaplan-Meier plot representing association between the age at CRC diagnosis and median separation of *AP2M1* expression in *APC*^*mut–*^ and *APC*^*mut+*^ CRCs from the TCGA dataset. (*D*) Flowchart of two molecular mechanisms that may be involved in the development of *APC*^*mut–*^ CRC.

## DISCUSSION

In the present study, we found potentiation of WNT signaling in *APC*^*mut–*^ CRCs was observed to occur through two molecular mechanisms represented in the CL1 and CL2 subgroups (Figure 5D). CL2 CRCs exhibited enhanced sensitivity to extracellular WNT signaling. Enhanced sensitivity was characterized by a striking downregulation of *RNF43* gene expression. Mechanistically, enhanced sensitivity occurred through several different molecular aberrations, including mutations of *RNF43, BRAF, PTPRK-RPSO3* gene fusion, DNA methylation and down-regulation of *AXIN2*. We developed the WNT_LS_ score to estimate the activation potential of extracellular WNT signaling in a CRC sample and found a high WNT_LS_ score was associated with CL2 *APC*^*mut–*^ CRCs. These results were validated in two independent datasets.

In addition to enhanced sensitivity to extracellular WNT signaling, CL2 *APC*^*mut–*^ CRCs exhibited more immune infiltrations compared to *APC*^*mut+*^ and to CL1 *APC*^*mut–*^ CRCs. Immune cell infiltration was notably higher in *APC*^*mut–*^ CRCs that had *RNF43* or *BRAF* mutations. M2 macrophages had the strongest association with potentiating WNT signaling in CRCs, due to their statistically significant correlation with *RSPO3* expression and *AXIN2* DNA methylation. Previous studies have shown that macrophages also have the capability to stimulate WNT signaling in response to tissue damage^36, 37^. The association of CL2 *APC*^*mut–*^ CRCs with M2 macrophages suggests the etiology of this cancer sub-type is tied to chronic tissue stress and inflammation that eventually favors a clone with hypersensitivity to WNT. We suggest that CL2 *APC*^*mut–*^ CRCs may be sensitive to porcupine inhibitors or anti-WNT/anti-DKK1 biologics.

CL1 *APC*^*mut–*^ CRCs were associated with low WNT_LS_ score and may be dependent on enhanced mitochondrial activation. *APC*^*mut–*^ CRC cell lines from the DepMap database had a strong dependency on mitochondrial activation relative to *APC*^*mut+*^ CRC cell lines. We are cautious in interpreting these data, because the observed effectiveness of mitochondria disruption of the *APC*^*mut–*^ CRC cell lines may be due to the absence of immune cells *in vitro*. One potential reason why some *APC*^*mut–*^ CRCs become dependent on enhanced mitochondrial activation is because mitochondria can stimulate the WNT pathway independently of WNT ligands^38^. Moreover, intestinal epithelial cell-specific knockout of TFAM, a transcription factor required for replication of mitochondria DNA, drastically reduced tumor formation in *APC*^*min/+*^ mouse models^39^. Therefore, we suggest that enhanced activation of mitochondria is a second, independent mechanism by which *APC*^*mut–*^ CRCs exploit WNT signaling in tumor progression. These findings also suggest that mitochondria inhibitors may be a promising therapeutic option for CL1 *APC*^*mut–*^ CRCs.

Based on a global gene expression analysis, we found that earlier-onset *APC*^*mut–*^ CRCs had increased expression of *AP2M1. AP2M1* plays an important role in clathrin-mediated endocytosis^40^. A recent study showed that when insulin binds to an insulin receptor, *IRS1* and *IRS2* recruit *AP2M1* to initiate insulin receptor endocytosis^41^. Thus, an increase of *AP2M1* may suggest increased insulin signaling. Importantly, insulin can activate both the PI3K pathway and the MAPK pathway, which may have roles in the observed enhanced mitochondria activation and increased immune infiltration subtypes of *APC*^*mut–*^ CRCs^42-44^. Other studies have found that individuals with type two diabetes are at a greater risk for early-onset CRC^45, 46^.

Early-onset CRC is a rapidly advancing public health emergency, and it is associated with a lack of mutation in APC. Our comprehensive genomic analysis has uncovered two classes of *APC*^*mut–*^ CRCs, one which potentiates WNT signaling through sensitivity to extracellular signaling, and the other which exhibits mitochondrial activation. Future research should test the effect of anti-WNT biologics and mitochondrial inhibitors in organoid models and in vivo.

## ADDITIONAL INFORMATION

## Acknowledgements

Special thanks to Ellen Kaye and the Maltz Foundation for their generous support of University of Arizona Cancer Biology GIDP student AG.

## Authors’ contributions

Study concept and design: AG, RMX, XL, NE, MP. Acquisition of data: AG. Analysis and interpretation of data: AG, RMX, VN, JL, CT, BS, XL, NE, MP. Drafting of manuscript: AG, MP. Critical revision of the manuscript for intellectual content: AG, RMX, CT, BS, XL, NE, MP. Statistical analysis: AG, MP. Obtained funding: XL, NE.

## Data availability

Data analyzed in this study can be found in the Genomic Data Commons (https://gdc.cancer.gov); Gene Expression Omnibus (GSE35896; https://www.ncbi.nlm.nih.gov/geo/); cBioPortal (CPTAC-2; https://www.cbioportal.org/); and DepMap (https://depmap.org/portal/). All analytic methods and study materials are available to other researchers through supplemental materials and in the methods section.

## SUPPLEMENTARY FIGURE LEGENDS

**Supplementary Figure 1**. (*A*) The fraction of *BRAF* mutations in *APC*^*mut–*^ and *APC*^*mut+*^ CRCs from the GSE35896 dataset. (*B*) The fraction of *RNF43, ZAP70, ERBB3, BRAF* and *ADGRL1* mutations in *APC*^*mut–*^ and *APC*^*mut+*^ CRCs from the CPTAC-2 dataset.

**Supplementary Figure 2**. Comparison of *ZNRF3* expression (*A*) and *RSPO3* expression (*B*) between *APC*^*mut–*^ CRCs, *APC*^*mut+*^ CRCs, and normal colon samples from the TCGA, GSE35896, and CPTAC-2 datasets. A two-sample t-test with a two-tailed p-value was used to determine statistical significance.

**Supplementary Figure 3**. Differentially expressed genes (adj p < 0.05) between *APC*^*mut–*^ CRCs and normal colon samples (*A*) and *APC*^*mut+*^ CRCs and normal samples (*B*) were mapped onto the KEGG canonical WNT signaling pathway. Blue labeled genes represent downregulation relative to normal samples, while red labeled genes represent upregulation relative to normal colon samples.

**Supplementary Figure 4**. Unsupervised clustering of *APC*^*mut–*^ CRCs from the GSE35896 dataset (*A*) and the CPTAC-2 dataset (*B*) using differentially expressed genes identified from comparing *APC*^*mut–*^ and *APC*^*mut+*^ CRCs from TCGA. Scatter plots for the GSE35896 dataset (*D*) and the CPTAC-2 dataset (*E*) showing the estimation of activation potential of extracellular WNT signaling for *APC*^*mut–*^ CL1, *APC*^*mut–*^ CL2, and *APC*^*mut+*^ CRCs. The y-axis represents a sample’s apparent sensitivity to extracellular WNT signaling using the WNT ligand sensitivity score. The x-axis represents a sample’s WNT stimulation potential by quantifying each sample’s maximum WNT ligand expression.

**Supplementary Figure 5**. Enhanced mitochondrial activation in CL1 *APC*^*mut–*^ CRCs. *(A)* A volcano plot representing differential expression analysis between CL1 *APC*^*mut–*^ and *APC*^*mut+*^ CRCs from the TCGA dataset. Blue points were downregulated in CL1 *APC*^*mut–*^ CRCs and red points upregulated. (*B*) A volcano plot representing differential expression analysis between CL2 *APC*^*mut–*^ and *APC*^*mut+*^ CRCs from the TCGA dataset. Blue points were downregulated in CL2 *APC*^*mut–*^ CRCs and red points upregulated. *(C)* An enrichment plot of the most significant upregulated GO term from GSEA analysis between CL1 *APC*^*mut–*^ and *APC*^*mut+*^ CRCs from the

TCGA dataset. (*D*) Unsupervised cluster analysis of *APC*^*mut–*^ CRCs from the TCGA dataset using genes associated with the biological process GO term Oxidative Phosphorylation. Shown are each *APC*^*mut–*^ CRC’s CIBERSORTx absolute score (representing total number of estimated infiltrating immune cells), presence of a *PTPRK-RSPO3* fusion, *BRAF* or *RNF43* mutation and CL1 or CL2 cluster from Figure 2D. (*D*) Violin plots of CRISPR dependency scores of the mitochondria-related genes *MRPL40, MRPL13, NDUFB8, NUDFB3* in *APC*^*mut–*^ CRC cancer cell lines (n = 3) and *APC*^*mut+*^ CRC cancer cell lines (n = 16). A Welch’s two-sample t-test with a “greater than” alternative hypothesis was used to test for statistical significance.

**Supplementary Figure 6**. Unsupervised cluster analysis of *APC*^*mut–*^ CRCs from the GSE35896 dataset *(A)* and the CPTAC-2 dataset *(B)* using genes associated with the biological process GO term Oxidative Phosphorylation. Shown are each *APC*^*mut–*^ CRC’s CIBERSORTx absolute score representing total number of estimated infiltrating immune cells.

