## Supplementary Figures for "Molecular Drivers of Tumor Progression in Microsatellite Stable *APC* Mutation-Negative Colorectal Cancers"

### Supplementary Figure 1

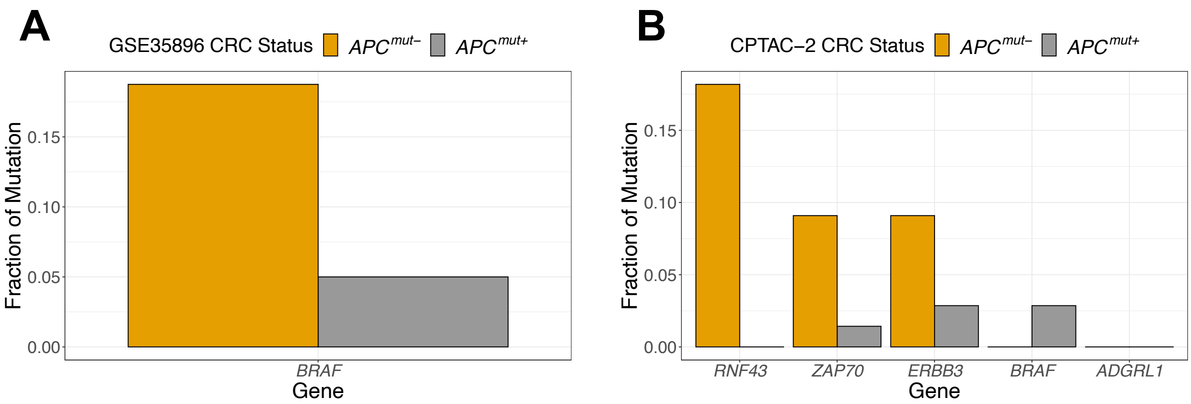

### Supplementary Figure 2

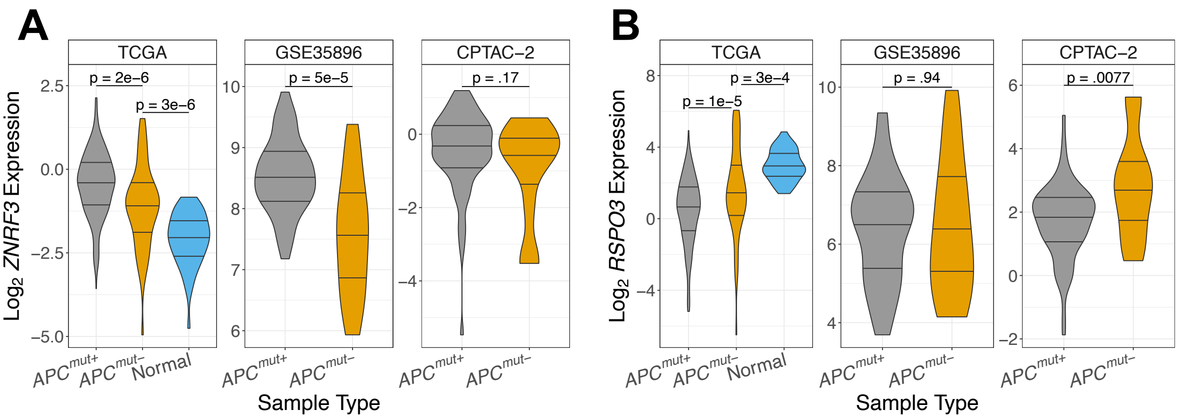

### Supplementary Figure 3

## A

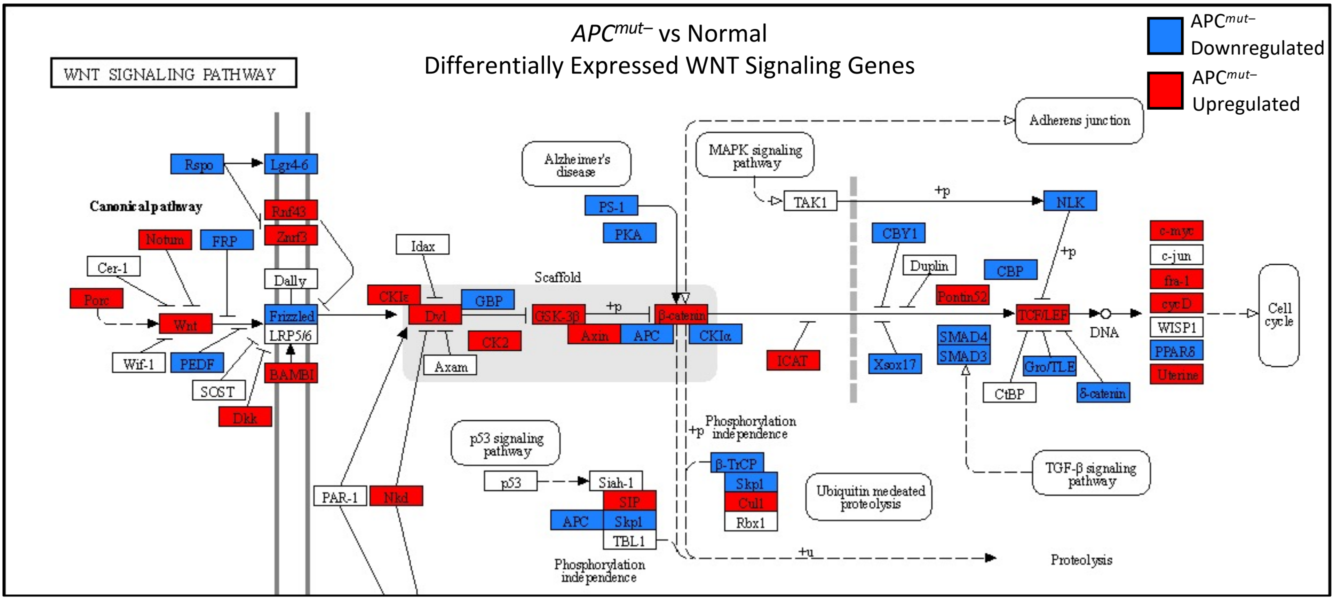


## B

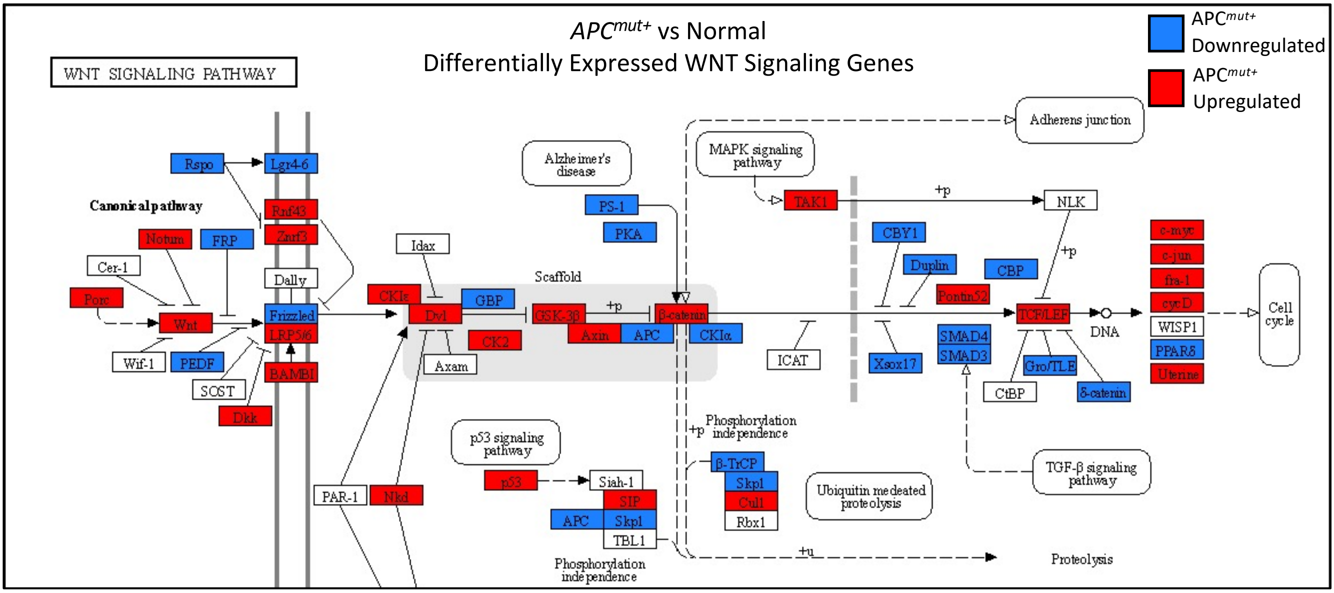

### Supplementary Figure 4

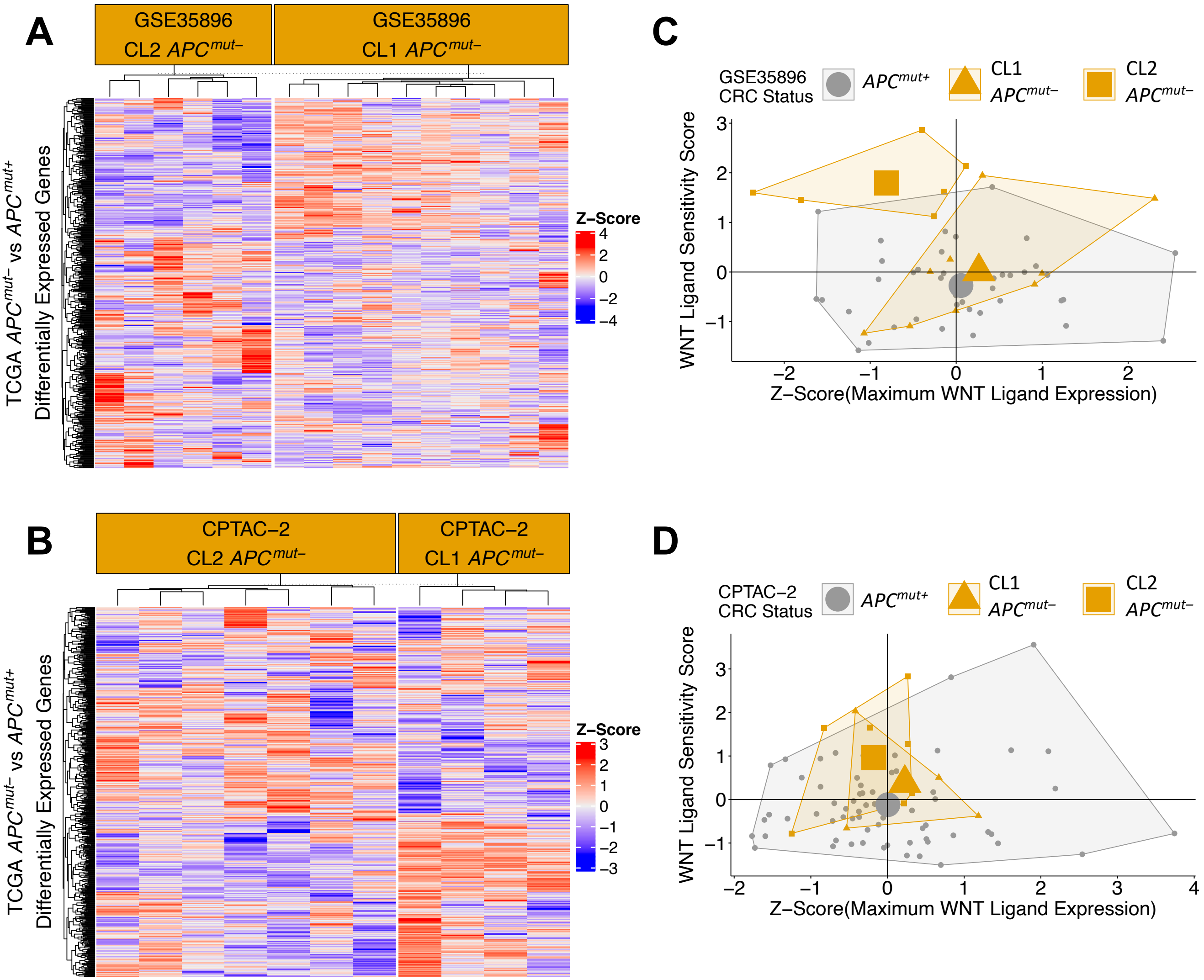

### Supplementary Figure 5

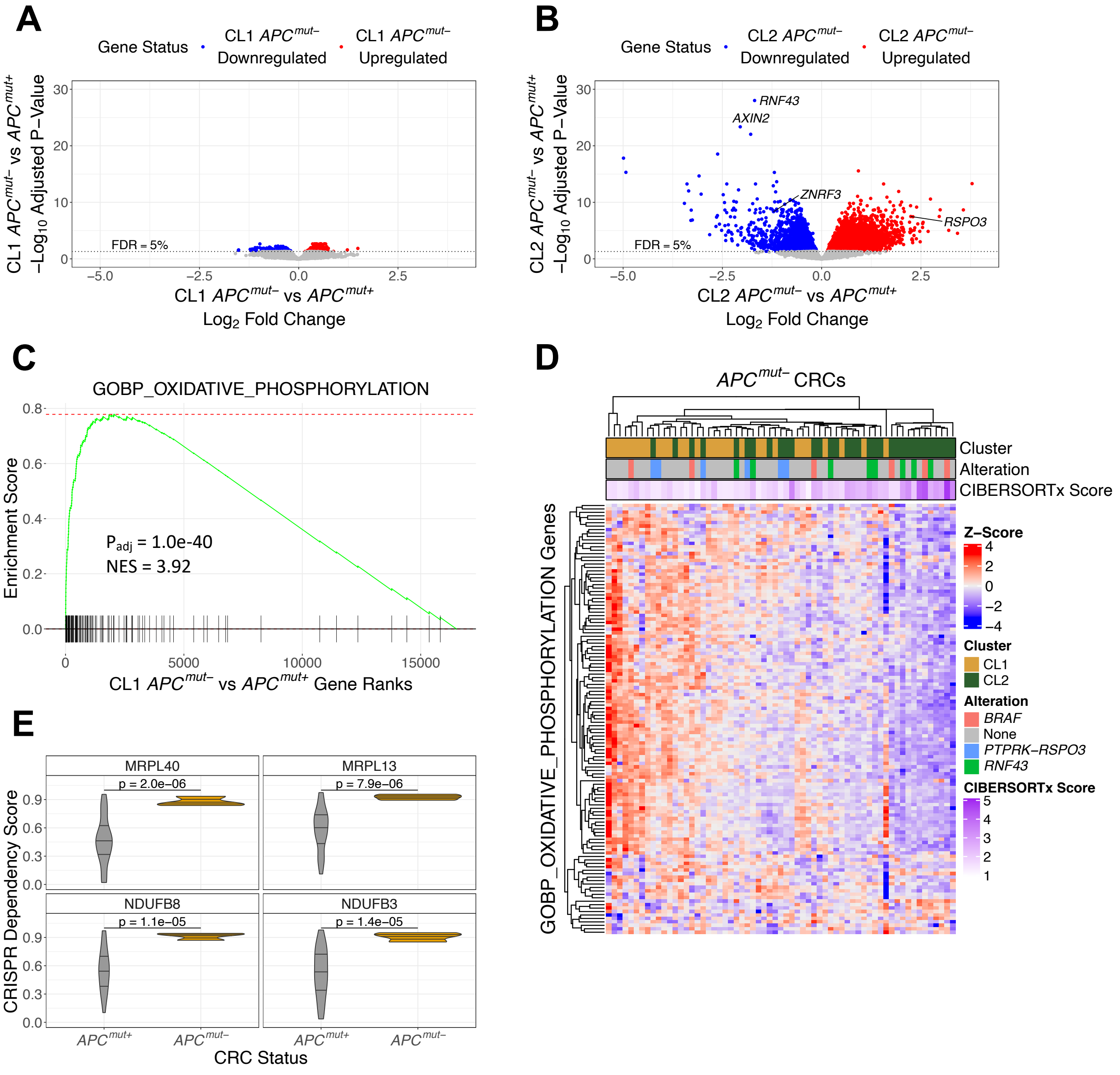

### Supplementary Figure 6

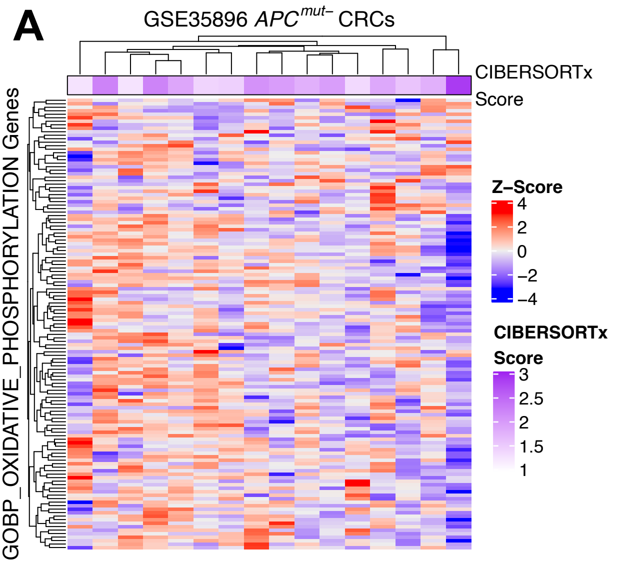
