## Supplementary Methods for "Molecular Drivers of Tumor Progression in Microsatellite Stable *APC* Mutation-Negative Colorectal Cancers"

***Discovery dataset: The Cancer Genome Atlas (TCGA)***

Genomic alterations. Colon adenocarcinoma (COAD) and rectal adenocarcinoma (READ) Mutect2 MAF files were downloaded from the Genomics Data Commons data portal using the R package TCGAbiolinks^1, 2^. The “pipeline” parameter in the GDCquery_Maf function was set to “mutect”. Mutations that were not labeled as PASS in the MAF file “FILTER” column and labeled as Silent in the “Variant_Classification” column were discarded from our study. Copy number variation data were extracted from the file “all_thresholded.by_genes.txt” in the TCGA FireBrowse data portal. Genes that were labeled as -2 were considered deep deletions while genes labeled 2 were considered amplifications. The TumorFusions study was used to call gene fusions in TCGA CRC samples^3^.

MSS and hypermutation. To determine which TCGA samples were microsatellite stable, we used the “COADREAD.clin.merged.txt” file from the FireBrowse data portal. Samples that were labeled as “MSI-high” were discarded from our study, except for the two samples TCGA-DC-6154 and TCGA-CK-6747, which had only 118 and 191 mutations, respectively. Next, we removed hypermutated samples. Based on the distribution of total mutations in TCGA MSS CRCs, we selected a hard cutoff of 700 total mutations to define hypermutation. In addition, we removed seven samples that had a non-passing *APC* mutation call. After these initial filtration steps, we were left with 425 CRC samples. In this study, CRC samples were classified as *APC^mut–^* if they lacked a non-silent mutation in *APC* (n = 75), did not have a deep deletion of *APC* (n = 411), and lacked a mutation in *CTNNB1* (n = 411). Using these criteria, we identified 63 *APC^mut-^* samples and 362 *APC^mut+^* samples.

Transcriptomics. We obtained HTSeq count files from the Genomics Data Commons^4^ and used the R packages edgeR and limma for normalization^5, 6^. Genes that had a CPM of less than 1 in at least 26 samples (half of the size of the smallest sample group—normal colon) were discarded from the study. To evaluate the expression data for batch effects, we used the R package MBatch and information from <https://bioinformatics.mdanderson.org/BEV/QF/>. Based on the recommended MBatch Overall-DSC threshold of 0.5, we did not identify a batch effect in any of the tested batch types: Batchld (0.48), Plateld (0.45), Platform (0.18), ShipDate (0.42), or tissue source site (0.38).

***Validation datasets: GSE35896 and CPTAC-2***

Genomic alterations. The GSE3896 dataset was downloaded from the Colorectal Cancer Subtyping Consortium on synapse.org^7, 8^. Consensus molecular subtype (CMS) and CpG island methylator phenotype (CIMP) information were extracted from the “clinical_molecular_public_all.txt” file. Mutation, CNV and microsatellite information for CPTAC-2 was downloaded from cBioPortal^9, 10^.

MSS and hypermutation. MSS and hypermutation in the GSE35896 and CPTAC-2 datasets were defined as similarly to the TCGA analysis as possible. Since the GSE35896 dataset does not have whole exome sequencing or copy umber variation data, we could not filter for hypermutation, *APC* deep deletion or *CTNNB1* mutations. Based on the data available, we classified 16 out of 56 MSS CRC samples as *APC^mut–^*. The CPTAC-2 dataset has microsatellite instability, whole exome sequencing and copy number variation data. In the CPTAC-2 dataset, we identified 81 CRC samples that were MSS and non-hypermutated. Out of these 81 CRC samples, 13 did not have an *APC* mutation, 78 did not have an *APC* deep deletion and 76 were *CTNNB1* mutation negative. Applying these criteria resulted in 11 *APC^mut–^* CRCs and 70 *APC^mut+^* CRCs.

Transcriptomics. For GSE35896, RMA normalized microarray data was downloaded from synapse.org. For CPTAC-2, we used the same RNA-seq preprocessing pipeline as for TCGA, with one alteration: genes that had a CPM of less than 1 in at least 5 samples were discarded from the study.

***Comparing APC^mut+^ and APC^mut–^ CRCs***

Genomic alterations. We used Fisher’s exact test to identify genomic alterations that were overrepresented in *APC^mut–^* CRCs relative to *APC^mut+^* CRCs. The R package “cometExactTest” was used to test for mutual exclusivity among pairs of genomic alterations in the *APC^mut–^* CRCs^11^.

Transcriptomics analyses. We used the R package limma to identify differentially expressed genes between *APC^mut–^* and *APC^mut+^* CRCs. Genes with a P_adj_ < 0.05 (Benjamini-Hochberg adjustment) were considered to be differentially expressed. The web application PathView was used to map differentially expressed genes onto the KEGG WNT canonical signaling pathway, with the node sum parameter set to “max.abs”^12, 13^. Gene set enrichment analysis (GSEA) analysis was performed using the R package fgsea^14^. As input for GSEA, we used the c5.go.bp.v7.4.symbols.gmt.txt file downloaded from http://www.gsea-msigdb.org/gsea/msigdb/collections.jsp and ranked the genes by the t-statistic from our differential expression analysis between *APC^mut–^* CRCs and *APC^mut+^* CRCs^15^. The Cytoscape application EnrichmentMap was used to visualize GSEA GO term results in network form^16, 17^. Only GO terms with P_adj_ < 0.05 were used to generate clusters.

CIBERSORTx analyses. The CIBERSORTx web application was used to impute the fraction of immune cells within gene expression data from the TCGA, GSE35896 and CPTAC-2 datasets^18^. When executing CIBERSORTx, we used the LM22 signature matrix file, enabled batch correction, disabled quantile normalization, and ran in absolute mode.

***Estimating WNT signaling transduction competence***

We developed a scoring system to quantify the WNT signaling transduction competence of each CRC tumor based on the mRNA expression of FRIZZLED (FZD) receptors and WNT ligands. When comparing *APC^mut–^* CRCs and *APC^mut+^* CRCs, we found significant changes in the expression of *RNF43* and *ZNRF3,* which inhibit the transduction of extracellular WNT signaling, and in *RSPO3*, which activates extracellular WNT signaling in part by inhibiting *ZNRF3* and *RNF43*. We therefore defined the WNT ligand sensitivity (WNT_LS_) score as the normalized expression of *RSPO3* minus the sum of the normalized expression levels of *RNF43* and *ZNRF3*: WNT ligand sensitivity = Z(*RSPO3_mRNA-z_* – *(RNF43_mRNA-z_* + *ZNRF3_mRNA-z_)*), where the subscript mRNA-z denotes the z-score of the expression value computed across all samples (Normal, *APC^mut-^* and *APC^mut+^).* However, a low level of negative regulators may not signify a sensitivity to extracellular WNT signaling and could be just a marker of low overall WNT pathway activity. To distinguish these two possibilities, we also computed the maximum WNT ligand expression over all 12 WNT ligands in our dataset for each CRC sample. Tumors with a high WNT_LS_ score that results in high expression of genes that code for WNT ligands are considered to have high WNT signaling transduction competence.

***DNA methylation preprocessing and analyses***

IDAT files for the TCGA Illumina Human Methylation 450 arrays were retrieved using the GDCquery function from the R package TCGAbiolinks. Preprocessing and normalization were carried out with the function preprocessFunnorm from the R package minfi^19, 20^. The R package MBatch and information from https:/bioinformatics.mdanderson.org/BEV/QF/ were used to test for batch effects. Based on the MBatch Overall-DSC threshold of 0.5, we did not identify a batch effect in any of the tested batch-types: BatchId (0.36), PlateId (0.35), ShipDate (0.34), or tissue source site (0.36). The R package DMRcate was used to identify differentially methylated regions (DMRs) using default parameters^21^. We used the R package annotatr to annotate the identified DMRs^22^.

***DepMap data and analyses***

All DepMap data was obtained from <https://depmap.org/portal/download/>^23, 24^. Cancer cell lines that were labeled as “Colon/Colorectal Cancer” in the primary_disease column of the “sample_info.csv” file were selected for this study. The file “Chan_et_al_2019_Supplementary_Table_1.csv” was used to determine if a CRC cell line exhibited MSI. Based on the distribution of total mutations calculated from the “CCLE_mutations.csv” file, we used a hard cutoff of 800 mutations to distinguish hypermutated and CRC cell lines from non-hypermutated CRC cell lines. After filtering out hypermutated and MSI-high cell lines, we were left with 19 CRC cell lines that had CRISPR knockout data. To distinguish *APC^mut–^* CRC cell lines from *APC^mut+^* CRC cell lines, we followed the same protocol used for the TCGA dataset. Three CRC cell lines lacked deleterious nucleotide variants in *APC*, no CRC cell lines had an *APC* deep deletion based on values from the “CCLE_gene_cn.csv” file, and 18 CRC cell lines were negative for a mutation in *CTNNB1.* The intersection of these three criteria identified 16 *APC^mut+^* CRC cell lines and 3 *APC^mut–^* CRC cell lines. To rank CRISPR knockouts, we used the statistic from Welch’s two-sample t-test (alternative hypothesis “greater than”) applied to the dependency scores of *APC^mut–^* and *APC^mut+^* cell lines. Dependency scores were extracted from the file “Achilles_gene_dependency.csv” on the DepMap portal.
